# Clinicopathologic features of a feline SARS-CoV-2 infection model parallel acute COVID-19 in humans

**DOI:** 10.1101/2021.04.14.439863

**Authors:** Jennifer M. Rudd, Miruthula Tamil Selvan, Shannon Cowan, Yun-Fan Kao, Cecily C. Midkiff, Jerry W. Ritchey, Craig A. Miller

## Abstract

The emergence and ensuing dominance of COVID-19 on the world stage has emphasized the urgency of efficient animal models for the development of therapeutics and assessment of immune responses to SARS-CoV-2 infection. Shortcomings of current animal models for SARS-CoV-2 include limited lower respiratory disease, divergence from clinical COVID-19 disease, and requirements for host genetic modifications to permit infection. This study validates a feline model for SARS-CoV-2 infection that results in clinical disease and histopathologic lesions consistent with severe COVID-19 in humans. Intra-tracheal inoculation of concentrated SARS-CoV-2 caused infected cats to develop clinical disease consistent with that observed in the early exudative phase of COVID-19. A novel clinical scoring system for feline respiratory disease was developed and utilized, documenting a significant degree of lethargy, fever, dyspnea, and dry cough in infected cats. In addition, histopathologic pulmonary lesions such as diffuse alveolar damage, hyaline membrane formation, fibrin deposition, and proteinaceous exudates were observed due to SARS-CoV-2 infection, imitating lesions identified in people hospitalized with ARDS from COVID-19. A significant correlation exists between the degree of clinical disease identified in infected cats and pulmonary lesions. Viral loads and ACE2 expression were quantified in nasal turbinates, distal trachea, lung, and various other organs. Natural ACE2 expression, paired with clinicopathologic correlates between this feline model and human COVID-19, encourage use of this model for future translational studies.

**Author Summary:** Identifying an ideal animal model to study COVID-19 has been difficult, and current models come with challenges that restrict their potential in translational studies. Few lab animals naturally express the receptors necessary for viral infection (ACE2), and many fail to manifest clinical signs or pathology similar to that seen in humans. Other models (non-human primates, mink) are ideal for disease and transmission studies, but are restricted by cost, husbandry challenges, and scarce availability. Alternatively, cats naturally express ACE2 receptors, are naturally infected with SARS-CoV-2 and can transmit virus from cat-to-cat. Prior to this study, cats infected by oral/nasal routes have not displayed significant clinical disease or lung pathology. However, we demonstrate that direct inoculation of concentrated SARS-CoV-2 virus in the trachea of cats induces analogous clinical and pathologic features to hospitalized patients with acute COVID-19. Our results show that infected cats exhibit significant clinical signs during experimental infection (coughing, increased respiratory effort, lethargy, and fever) and exhibit extensive lung lesions that mimic severe COVID-19 pathology such as diffuse alveolar damage and hyaline membrane formation – highlighting the immeasurable potential for this feline model to address translational approaches for COVID-19 and to better understand the role of cats in transmission and disease.

## INTRODUCTION

Since the emergence of severe acute respiratory syndrome coronavirus-2 (SARS-CoV-2) in late 2019, Coronavirus Disease 2019 (COVID-19) has swept across the globe resulting in nearly 3 million deaths worldwide as of March 2021 (1). Although a wide range of clinical symptoms are reported, mortality of COVID-19 patients is closely correlated with progression of viral infection to severe lung disease (pneumonia) and respiratory failure due to acute respiratory distress syndrome (ARDS), which is further complicated by immune cell dyscrasias and hyperinflammation (cytokine storm) in critically ill patients (2-4). Features of pulmonary pathology that are hallmarks of severe COVID-19 (i.e., diffuse alveolar damage with hyaline membrane formation, type II pneumocyte hyperplasia, vascular thrombi, fibrin and serous exudation) have been difficult to reproduce in animal models, making it impossible to completely understand the pathophysiology of disease and to test efficacy of new therapeutics and vaccines (5, 6). Identification of a translational animal model that parallels clinical and pathologic features of disease in addition to route of infection, replication, and transmission kinetics is of paramount importance.

SARS-CoV-2 viral infection and replication within a host requires the presence and distribution of angiotensin-converting enzyme 2 (ACE2) receptors similar to humans (7). Natural SARS-CoV-2 infections in animals are documented to occur in a diverse range of species, including domestic and exotic cats, dogs, mink, and Golden Syrian hamsters (8-11), and this diverse host range is largely due to natural expression of ACE2 receptors and host tropism of this receptor with the S protein of SARS-CoV-2 (12, 13). Due to the natural availability of ACE2 receptors and confirmed host susceptibility and transmission (10, 14-17), domestic cats offer an exciting advantage as experimental models for SARS-CoV-2 infection (18, 19). Comorbidities that exacerbate COVID-19 disease, such as hypertension, diabetes, renal disease, and obesity, are readily adapted to feline models (20-25). Furthermore, establishing a SARS-CoV-2 infected feline model is prudent to better understand zoonotic transmission potential from domestic cats back to people in close contact.

Previous studies have successfully infected cats with SARS-CoV-2 via intra-nasal (1-3.05×10^5^ PFU) and/or intra-oral routes (5×10^5^ TCID_50_/ml) and have confirmed cat-to-cat transmission through both respiratory droplets and aerosolization (16, 26-28). However, these studies failed to produce clinical signs in infected cats, and evidence of lower respiratory pathology mirroring severe COVID-19 in humans was not observed (16, 26-28), potentially due to concentration of the viral inoculum and/or inoculation route. Interestingly, pulmonary disease with diffuse alveolar damage was previously documented in cats intra-tracheally infected with 1×10^6^ TCID_50_ SARS-CoV-1, which also resulted in efficient transmission of virus to uninfected animals (29, 30).

Based on outcomes of these former studies, we hypothesized that inoculation with a higher concentration of SARS-CoV-2 via the intra-tracheal route would result in pulmonary pathology and clinical disease in domestic cats similar to COVID-19 in human patients. The experiments reported in this study provide the first feline model of SARS-CoV-2 infection with significant lower respiratory disease that displays features of diffuse alveolar damage seen in the early exudative phase of human COVID-19. In addition, SARS-CoV-2 infected cats exhibited clinical signs of lower respiratory disease characterized by increased respiratory effort and coughing in addition to signs of systemic involvement such as pyrexia and lethargy. While the role of cats in zoonotic transmission is still under investigation, the applicability of a clinically significant SARS-CoV-2 feline model with pathological lesions that mirror severe COVID-19 is of high impact for future studies.

## RESULTS

### SARS-CoV-2 infected cats exhibit clinical signs of lower respiratory disease

In order to clinically assess the feline model in Animal Biosafety Level-3 conditions, a novel clinical scoring system for feline respiratory disease was developed by integrating features of previously utilized systems (31-33) (Table 1). Each cat was assigned a score from 0 to 2 for each of the following categories: body weight loss, activity levels, behavioral changes, body temperature, respiratory effort, ocular or nasal discharge, and coughing. Scores were then summated to assign an overall clinical score for each day.

**Table 1.**
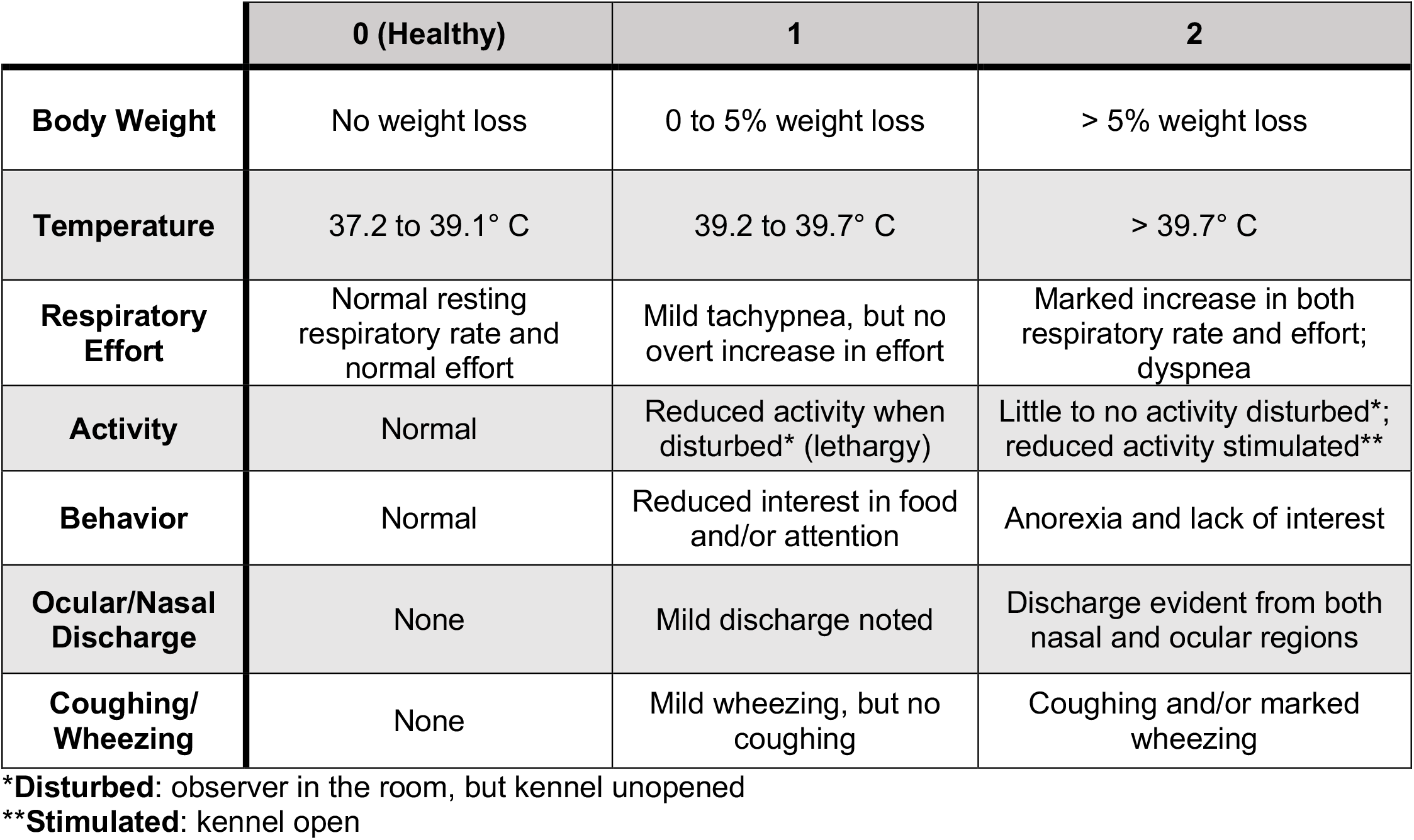
Clinical Scoring System for Feline Respiratory Disease. A scoring system was designed to assess clinical lower respiratory and systemic disease in the feline model. Each cat was scored daily at the same time point (morning) by a small animal clinician (JMR). Cats were assigned a score from 0 to 2 for each clinical parameter: body weight, temperature, respiratory effort, activity, behavior, ocular/nasal discharge, coughing/wheezing. The parameter scores were summed to assign an overall score per cat per day. Potential scores can range from 0 (healthy with no signs of disease) to 14 on any given day. Resting respiratory rate was considered normal if <36 breaths per minute. Marked increases in rate were >50 breaths per minute at rest. Temperatures were obtained via thermal microchips, and body weights were obtained last to limit stress affecting clinical scoring.

|  | 0 (Healthy) | 1 | 2 |
| --- | --- | --- | --- |
| **Body Weight** | No weight loss | 0 to 5% weight loss | > 5% weight loss |
| **Temperature** | 37.2 to 39.1° C | 39.2 to 39.7° C | > 39.7° C |
| **Respiratory Effort** | Normal resting respiratory rate and normal effort | Mild tachypnea, but no overt increase in effort | Marked increase in both respiratory rate and effort; dyspnea |
| **Activity** | Normal | Reduced activity when disturbed* (lethargy) | Little to no activity disturbed*; reduced activity stimulated** |
| **Behavior** | Normal | Reduced interest in food and/or attention | Anorexia and lack of interest |
| **Ocular/Nasal Discharge** | None | Mild discharge noted | Discharge evident from both nasal and ocular regions |
| **Coughing/Wheezing** | None | Mild wheezing, but no coughing | Coughing and/or marked wheezing |

\***Disturbed:** observer in the room, but kennel unopened
\*\***Stimulated:** kennel open

SARS-CoV-2-infected cats exhibited a significant increase in clinical disease scores starting on 4 days post-inoculation (dpi) and then on 5, 6, and 8 dpi when compared to sham-inoculated controls (Fig 1A). Clinical disease peaked on 4 dpi and continued through the study endpoint at day 8. The most prominent clinical signs noted were lethargy and increased respiratory effort, which were observed in 100% (12/12) of SARS-COV-2-infected cats during this study. Both lethargy and respiratory effort increase significantly between 3 and 4 dpi (*p*=0.0027; *p*=0.0027) and remained elevated with significantly higher scores through 8 dpi when compared with day 0 (Fig 1B). Coughing was noted in 4 of 12 infected cats with peak clinical signs occurring at 4 dpi. Pyrexia (temperature > 39.2°C) was documented in 8 of 12 SARS-CoV-2 infected cats over the course of the study, while 7 infected cats displayed altered behavior (reduced interest in food or attention) and 5 had measurable weight loss. No cats had ocular or nasal discharge (S1 Table). Sham-inoculated cats did not exhibit clinical signs except for one cat with mild weight loss on day 4.

**Fig 1.**
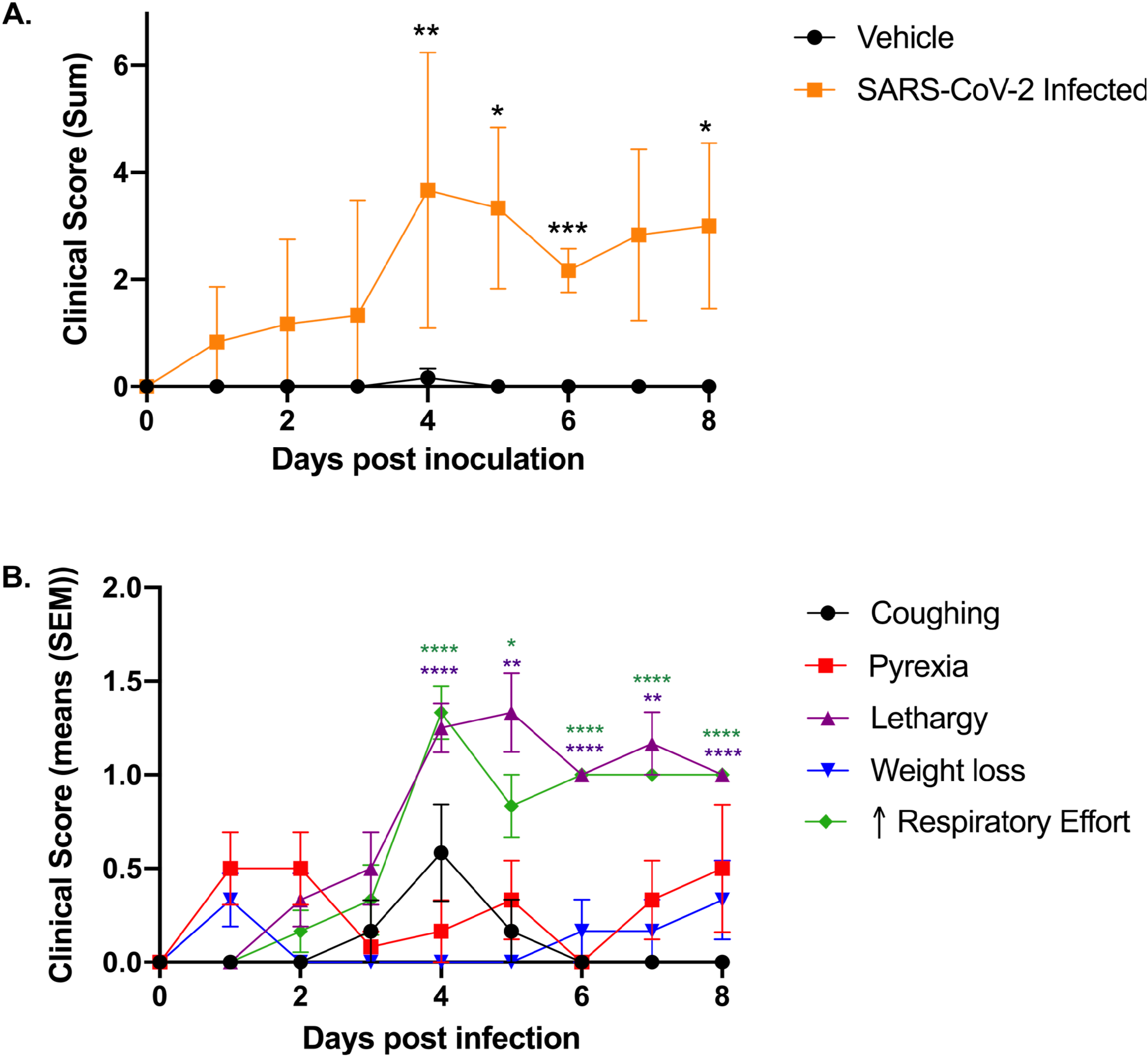
Intra-tracheal SARS-CoV-2 inoculation results in clinical disease. Clinical parameters were assessed using the feline respiratory disease clinical scoring system (see Table 1). **(A)** Clinical parameters summated to provide an overall clinical score per cat per day. Clinical disease severity peaked on 4 dpi and was significantly higher than sham-inoculated cats on 4 dpi (*p*=0.0054), 5 dpi (*p*=0.0257), 6 dpi (*p*=0.0004), and 8 dpi (*p*=0.0453). A noticeable trend in severity was also noted on 7 dpi as compared with sham-inoculated controls (*p*=0.0654). **(B)** Lethargy and increased respiratory effort were the most prominent clinical signs observed in SARS-CoV-2-infected cats; both of which were significantly increased between days 3 and 4 (*p*=0.0027; *p*=0.0027) and remained significantly elevated in infected cats after 4 dpi as compared to day 0. Coughing was most prominent on 4 dpi and was identified in 4/12 infected cats. Pyrexia was noted in 8/12 cats over the course of the study. Data are expressed as means ± SEM. Statistical comparisons made via mixed effects analysis. \**p*<0.05; \*\**p*<0.01; \*\*\**p*<0.001; \*\*\*\**p*<0.0001.

### Feline SARS-CoV-2 infection pathology mirrors acute COVID-19

Complete post-mortem evaluation was performed for all sham-inoculated control animals (n=6) and SARS-CoV-2-infected cats euthanized on day 4 (n=6) and day 8 (n=6) post-inoculation. Necropsy tissues from SARS-CoV-2-infected cats (including lung, trachea, nasal turbinates, and tracheobronchial lymph node (TBLN)) were grossly examined and compared to those from sham-inoculated cats (Fig 2 A-C). At 4 dpi, the lungs of SARS-CoV-2-infected cats were grossly heavy and wet, with large multifocal to coalescing regions of dark red consolidation that exuded a moderate amount of edema upon cut section (Fig 2 B). Gross lung lesions were similar at 8 dpi in SARS-CoV-2-infected cats, although the degree of pulmonary edema was moderately more pronounced (Fig 2 C). The TBLN of all SARS-CoV-2-infected cats were diffusely enlarged 4-5 times normal at both 4 dpi (n=6) and 8 dpi (n=6).

**Fig 2.**
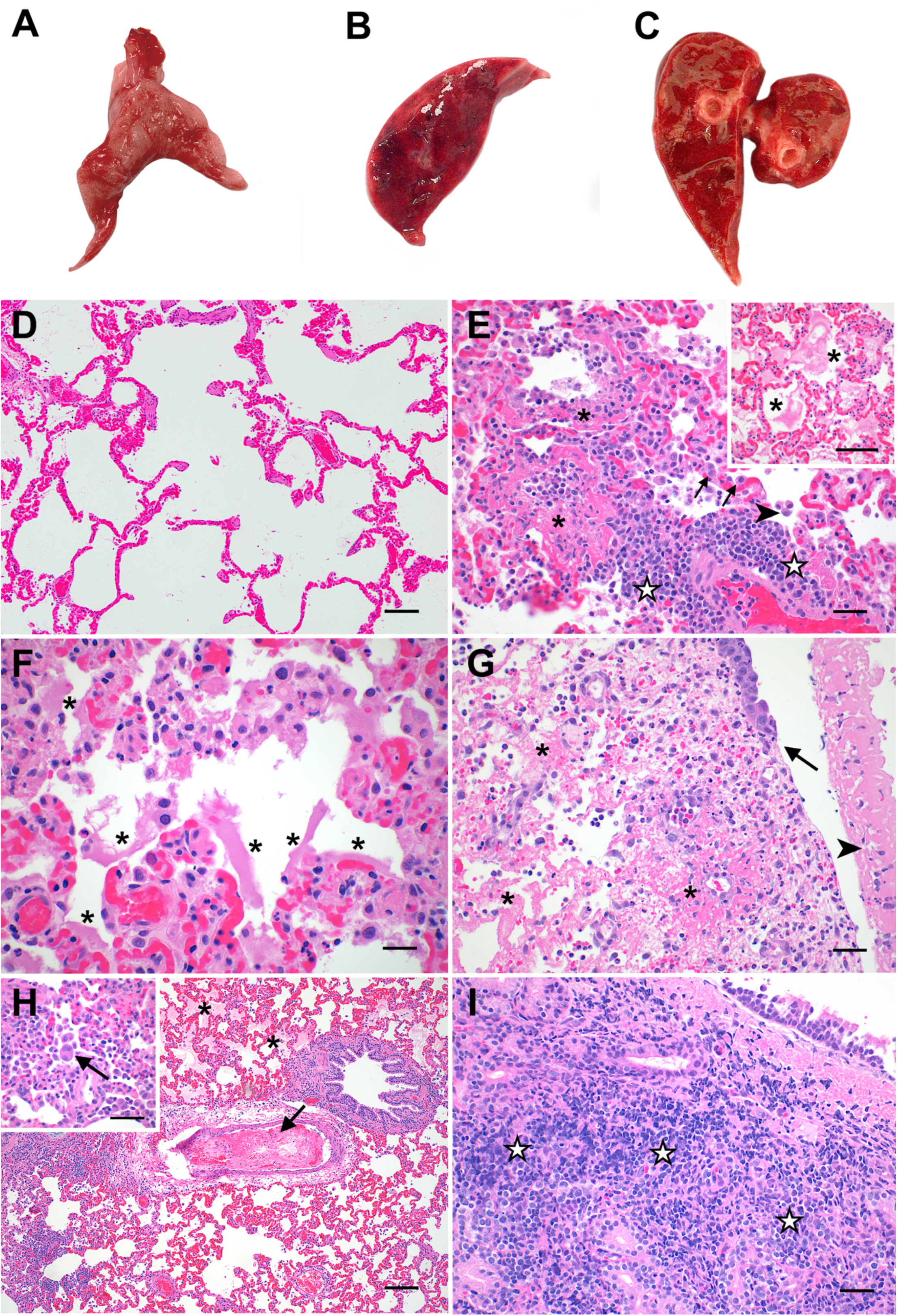
Pathologic features of acute SARS-CoV-2 infection in cats are analogous to the exudative phase of COVID-19. Compared to lungs from healthy sham-inoculated cats **(A)**, the lungs of SARS-CoV-2-infected cats were diffusely consolidated, dark red and edematous at both 4 dpi **(B)** and 8 dpi **(C)**. The lungs of healthy, uninfected cats **(D)** were histologically normal, with open alveoli and minimal atelectasis. At 4 dpi, the lungs of SARS-CoV-2-infected cats **(E)** exhibited discrete foci of alveolar inflammation and necrosis with fibrin deposition (**\***), increased alveolar macrophages (arrowhead), perivascular lymphocytes (**⋆**), and type II pneumocyte hyperplasia (arrows). The alveoli in these cats’ lungs were frequently filled with large amounts of edema and fibrin strands **(E** inset), and there were multifocal areas of hyaline membrane formation **(F)** (**\***). The distal trachea of 1 SARS-CoV-2 infected cat **(G)** was multifocally ulcerated at 4 dpi (arrow) with diphtheritic membrane formation (arrowhead) and multifocal areas of submucosal necrosis and fibrinoid vasculitis (**\***). At 8 dpi **(H)**, fibrinoid vasculitis, vascular thrombosis (arrow), and occasional syncytial cells **(H** inset) were observed in addition to the histopathologic changes described above. Tracheal lesions observed at 8 dpi **(I)** were characterized by varying degrees of lymphoplasmacytic, histiocytic, and neutrophilic inflammation with multifocal areas of submucosal necrosis. Magnification: (D, H) 10x, scale bar = 100µm; (E) 20x, scale bar = 50µm; (E inset, G, H inset, I) 40x, scale bar = 25µm; (F) 60x, scale bar = 17µm

Microscopic evaluation of selected necropsy tissues (lung, trachea, nasal turbinates, TBLN, and kidney) was performed for all study animals. Tissue sections from all sham-inoculated animals (n=6) were histologically unremarkable (Fig 2 D and S2 Table). In contrast, histopathologic features of feline SARS-CoV-2 infection exhibited striking similarities to documented pathologic features of the acute (exudative) and organizing phases of human COVID-19 (34-37). At 4 dpi, 100% (6/6) of SARS-COV-2-infected cats exhibited a significant degree of lung pathology (interaction, *p*<0.0001) and prominent histologic features consistent with diffuse alveolar damage (DAD) (Fig 2 E-F). Pulmonary edema (5/6 cats), multifocal alveolar damage and necrosis (5/6 cats), perivascular lymphocytic and neutrophilic infiltrates (6/6 cats), and increased intra-alveolar macrophages (5/6 cats) were significantly elevated in SARS-CoV-2-infected cats at 4 dpi (Supporting Information). These changes were occasionally accompanied by multifocal areas of hyaline membrane formation (3/6 cats), mild to moderate amounts of intra-alveolar fibrin (2/6 cats), type II pneumocyte hyperplasia (2/6 cats), and intra-alveolar syncytial cells (2/6 cats). One SARS-CoV-2 infected cat exhibited severe inflammation in the distal trachea at 4 dpi characterized by multifocal areas of submucosal necrosis and fibrinoid vasculitis with multifocal areas of mucosal ulceration and diphtheritic membrane formation (Fig 2 G).

Similar histologic features of DAD were also observed in the lungs of SARS-CoV-2-infected cats at 8 dpi, however, the overall pattern of lung injury appeared exhibited more prominent features of vascular injury compared to day 4 (Fig 2 H). A significant degree of pulmonary edema/exudate, perivascular inflammatory infiltration, and alveolar histiocytosis was present in 100% of SARS-CoV-2 animals (6/6 cats) at 8 dpi (Supporting Information). Alveolar damage and necrosis (4/6 cats) and intra-alveolar fibrin (3/6 cats) were also prominent features at this time point. Moreover, histologic evidence of fibrinoid vasculitis (2/6 cats) and vascular thrombosis (2/6 cats) was also observed at 8 dpi, in addition to occasional viral syncytia (1/6 cats) (Fig 2H). In 2 of these cats, the tracheal submucosa was multifocally expanded and effaced by moderate to severe lymphoplasmacytic, histiocytic, and neutrophilic inflammation with necrosis (Fig 2 I).

A positive linear correlation exists between peak clinical scores and histopathology scores of the lungs in SARS-CoV-2 infected cats (*p*=0.0002; R^2^=0.5884) indicating that severe clinical signs of disease correlate with pulmonary pathology (S1 Fig). Mild, multifocal lymphofollicular inflammation was observed in the nasal turbinates of all SARS-CoV-2-infected cats (6/6) at 4 dpi and in 4/6 cats at 8 dpi, with variable neutrophilic infiltration (Supporting Information). All SARS-CoV-2-infected animals (12/12) exhibited mildly increased lymphoid hyperplasia in TBLN at 4 and 8 dpi characterized by increased medullary cords and extranodal proliferations (Supporting Information), but overall changes were not statistically significant. No significant histopathologic findings were observed in renal tissues at either time point. Fluorescent immunohistochemistry was performed to detect SARS-CoV-2 positive cells in lung and TBLN of 2 SARS-CoV-2 infected cats (n=1 at 4 dpi, n=1 at 8 dpi). At both time points, low numbers of mononuclear cells positive for SARS-CoV-2 nucleoprotein were detected within the medulla of the TBLN, however, no positive cells were observed in lungs of these animals. (Fig 3).

**Fig 3.**
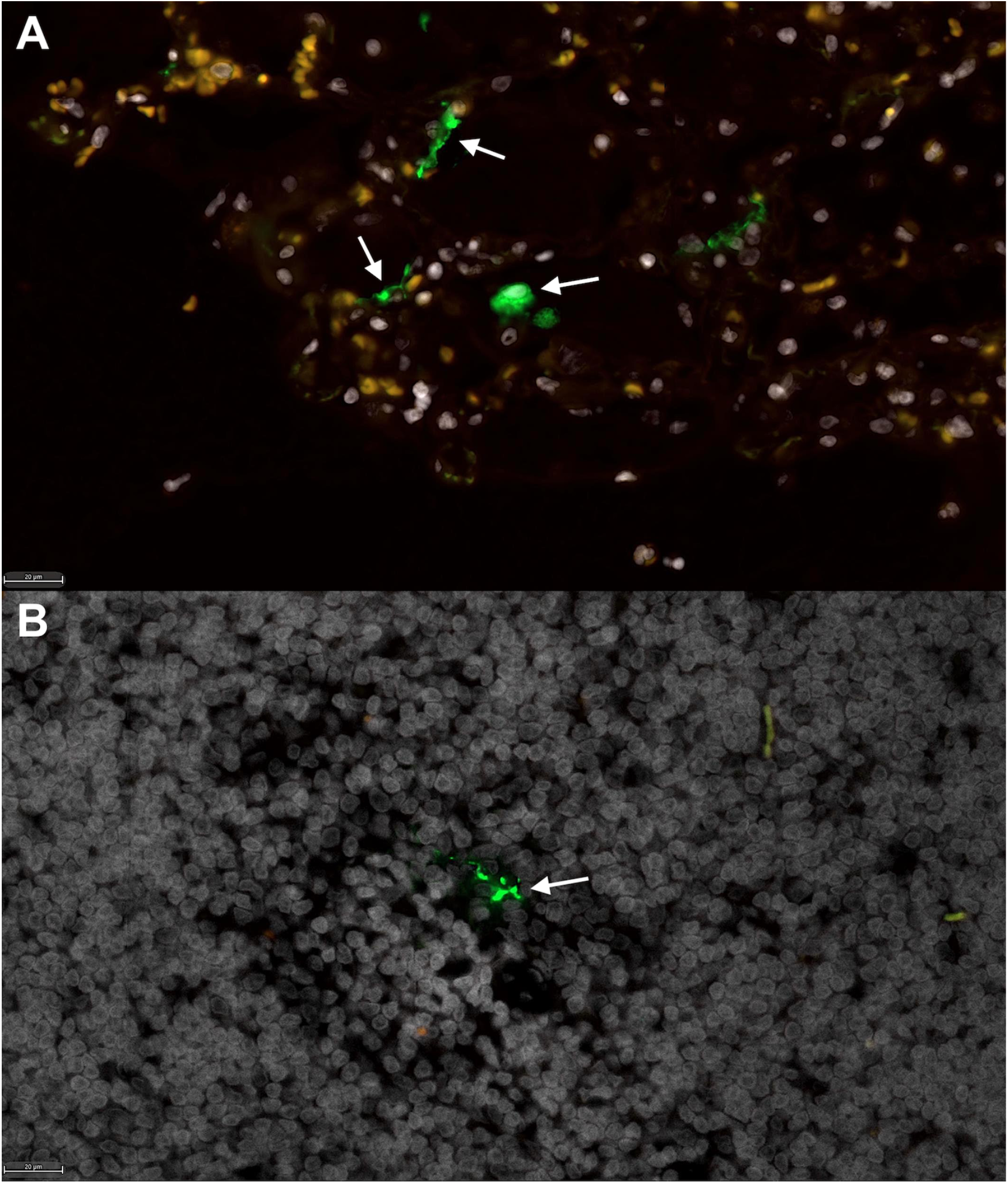
Fluorescent immunohistochemistry for SARS-CoV-2 nucleoprotein identifies mononuclear cells in tracheobronchial lymph node of intratracheally-infected cats. Low numbers of SARS-CoV-2 positive cells (green, white arrows) are detected in **(A)** positive control tissue (lung) from an African Green Monkey infected with SARS-CoV-2 (51), and within **(B)** mononuclear cells in the TB LN of SARS-CoV-2 infected cats (green, white arrow). White = DAPI/nuclei; green = CoV-2. Magnification (A-B) 40x, scale bar = 20 µm.

### ACE2 expression and viral RNA in feline tissues during SARS-CoV-2 infection

SARS-CoV-2 viral RNA and fACE2 RNA expression was quantified in the nasal turbinates, TBLN, distal trachea, kidneys and lungs of all SARS-CoV-2-infected cats (n=12) and sham-inoculated controls (n=6) using ddPCR (Fig 4 and S3 Table). Viral RNA was detected in 100% of tissues collected on 4 dpi from SARS-CoV-2-infected cats (Fig 4 A). At 8 dpi, viral RNA was also detectable in the lung, TBLN, and kidney tissues of all (6/6) infected cats, and in 5/6 cats in the nasal turbinates and 5/6 cats in the distal trachea. No SARS-CoV-2 viral RNA was detected in tissues collected from sham-inoculated cats at either time point (S3 Table). SARS-CoV-2 viral RNA copies were elevated in the TBLN at 8 dpi compared with day 4 samples, although this trend was not significant (*p*=0.0567). In contrast, SARS-CoV-2 viral load in lung samples was significantly lower at 8 dpi than at 4 dpi (*p*=0.0007) (Figure 4 A). A positive linear correlation was observed between SARS-CoV-2 RNA in the lung and pulmonary histopathology scores of SARS-CoV-2 infected cats (*p*=0.0183; R^2^=0. 3012) (S1 Fig). SARS-CoV-2 RNA was not reliably detected in nasal swabs or plasma of infected cats at either time point.

**Fig 4.**
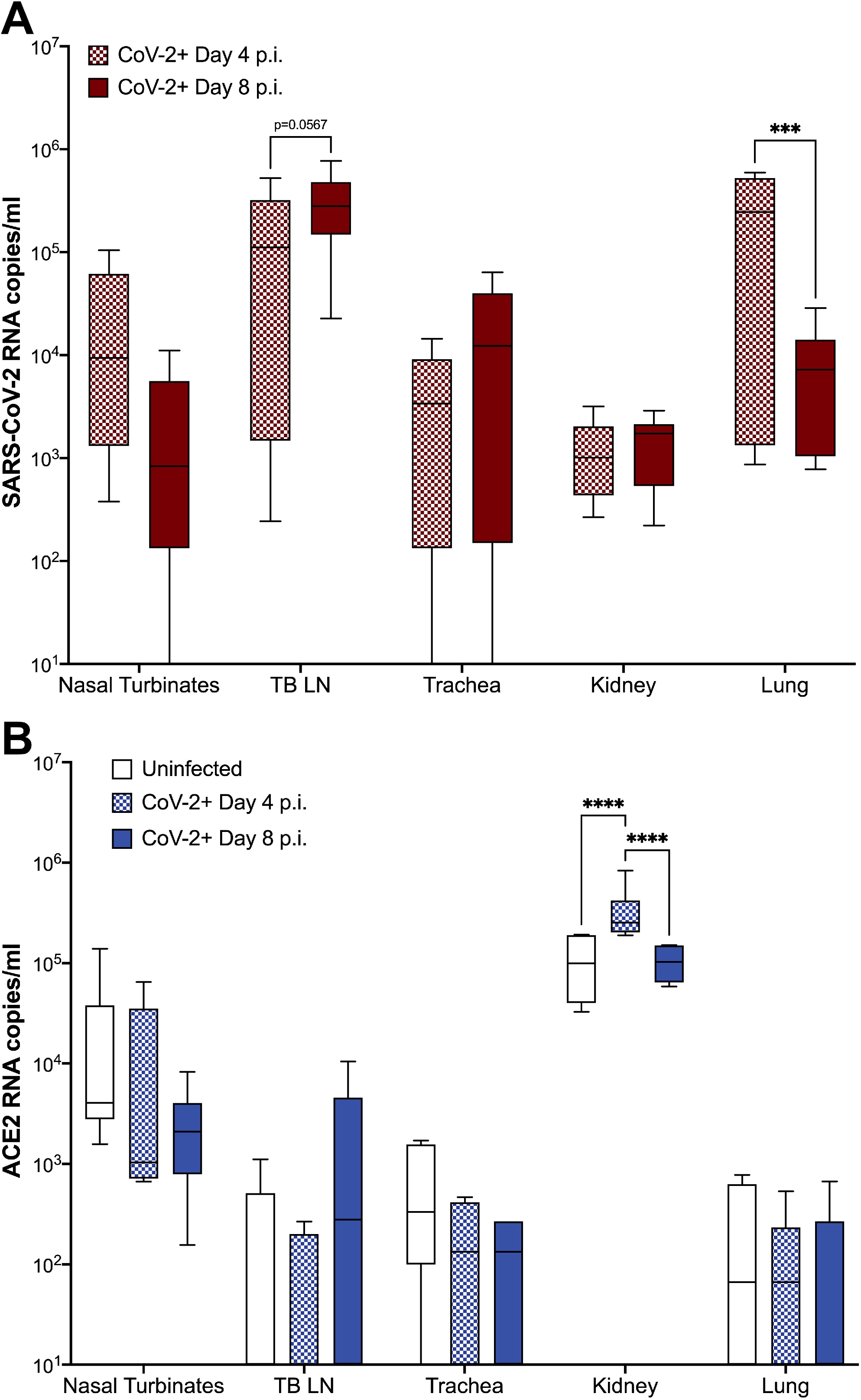
fACE2 RNA and SARS-CoV-2 viral RNA quantification in feline tissues. Extraction of SARS-CoV-2 and ACE2 RNA was performed as described from tissues samples collected on either 4 or 8 dpi. Tissue samples included nasal turbinates, tracheobronchial lymph node (TB LN), trachea, kidney, and lung. **(A)** SARS-CoV-2 RNA copies were detected in all tissues collected from cats inoculated with SARS-CoV-2. No viral RNA was detected in tissues from sham-inoculated cats. In SARS-CoV-2-infected cats, viral RNA copies were slightly increased in the TB LN between 4 to 8 dpi (*p*=0.0567) while viral RNA load in the lungs significantly decreased over the same period (*p*=0.0007). **(B)** fACE2 receptor RNA is significantly increased in the kidney of SARS-CoV-2 infected cats at 4 dpi compared to sham-inoculated controls (*p*<0.0001) SARS-CoV-3-infected cats at 8 dpi (*p*<0.0001). Data are expressed as means ± SEM. n=6 cats per group. Statistical comparisons made via two-way ANOVA. \*\*\**p*<0.001; \*\*\*\**p*<0.0001.

In sham-inoculated cats, Kruskal Wallis test revealed that fACE2 RNA in the nasal turbinates was significantly higher than in the lung (*p*=0.0093) and TBLN (*p*=0.0049). fACE2 RNA was also higher in the kidney when compared to lung (*p*=0.0003), trachea (*p*=0.0034), and TBLN (*p*=0.0001). These findings were similar in SARS-CoV-2-infected animals at 4 and 8 dpi, with fACE2 RNA levels being significantly higher in the nasal turbinates and kidney versus other tissues (*p*<0.05) (S4 Table). Overall, fACE2 RNA in the kidney was significantly increased in SARS-CoV-2 infected cats on 4 dpi when compared to both sham-inoculated controls, as well as SARS-CoV-2 infected cats on 8 dpi (ANOVA, *p*<0.0001) (Figure 4 B). No other significant changes in ACE2 RNA were observed over time.

## DISCUSSION

The potential of this feline model for future evaluation of COVID-19 is extensive. Challenges with earlier feline models of SARS-CoV-2 infection included a lack of clinical disease and/or pathology of the lower respiratory tract that resembles lesions seen in patients with COVID-19. The differences in clinical presentation between previous feline models and the model described here are likely attributed to modifications in routes and concentration of inoculation. In this study, SARS-CoV-2 was inoculated through an intra-tracheal route and at a higher concentration than previously reported (16, 26, 27). Route of inoculation is an important consideration when establishing an animal model for disease, and previous studies have exhibited marked differences in primary disease severity and distribution based on route of inoculation (38, 39).

While previous feline models offer value for study of asymptomatic infections, viral shedding, and transmission of SARS-CoV-2, cats infected through an intra-tracheal route exhibit clinical disease that aligns with that seen in early phases of acute COVID-19. Clinical assessment of infectious lower respiratory disease in a cat can be challenging, and it is not uncommon for cats with confirmed histologic infectious pneumonia to have limited clinical respiratory signs (40). Therefore, the clinical signs of respiratory disease induced in this model are highly significant. A novel clinical scoring system was designed that could be applied in the Animal Biosafety Level-3 facility to carefully assess for clinical disease. Interestingly, the disease noted in the SARS-CoV-2 infected cats was similar to that described in hospitalized patients with COVID-19. Clinical disease of hospitalized human COVID-19 patients is characterized by fever (70-90%), dry cough (60-86%), shortness of breath (53-80%), and fatigue (38%) (41) while predominant clinical sings in SARS-CoV-2 infected cats consisted of fever, cough, lethargy and increased respiratory effort, with lethargy and increased respiratory effort being the most notable clinical signs (Fig 1).

In addition to clinical signs of lower respiratory and systemic disease, SARS-CoV-2 infected cats also exhibited conspicuous pulmonary lesions of diffuse alveolar damage (DAD) by 4 dpi, and additional evidence of vascular damage by 8 dpi (Fig 2). Specific histopathological lesions align closely with those reported in human COVID-19 patients (34-36, 42-44), including DAD resulting in hyaline membrane formation, type II pneumocyte hyperplasia, occasional intra-alveolar syncytial cells, and the development of fibrinous exudate and vascular thrombi. To the author’s knowledge, this is the first report of hyaline membrane formation and type II pneumocyte hyperplasia in feline SARS-CoV-2 infection. Peak clinical disease scores positively correlated with severity of histologic lesions in the lungs (S1 Fig), which further support that cats with marked pulmonary histologic damage also had more severe clinical signs of disease.

Surprisingly, intra-tracheal inoculation of SARS-CoV-2 did not produce high viral RNA loads in the lungs as compared with other studies in which the inoculate was delivered via the intranasal route (26). However, despite by-passing the upper airway, virus was still detected in the nasal turbinates by 4 and 8 dpi, suggesting the virus may utilize the mucociliary escalator to travel up the respiratory tree and establish infection intra-nasally even without intra-nasal inoculation. Although seemingly lower quantities of SARS-CoV-2 RNA were recovered from lungs of intra-tracheally inoculated cats, the damage to lung tissues was highly evident, indicating that extensive pulmonary damage will occur even without high levels of viral replication within the pulmonary tissue at 4 and 8 dpi. Viral migration from lung to the TBLN occurred quickly (by 4 dpi) and this TBLN involvement is a novel finding to the feline model of SARS-CoV-2, including detection of viral antigen within the TBLN via fluorescent immunohistochemistry.

Similar to humans, fACE2 RNA expression varied by tissue location, but were relatively low in the lungs of both infected and uninfected cats. It is important to note that RNA measurements indicate an upregulation or downregulation of production of proteins, but do not necessarily indicate an absolute number of receptors available. However, it is possible that inefficient replication and rapid clearance of SARS-CoV-2 in the lungs is related to lower expression of ACE2 receptors as compared with nasal turbinate ACE2. Histopathology shows that cells regularly expressing ACE2 are damaged in the lung and this viral-induced pulmonary epithelial pathology may contribute further. ACE2 RNA copies in the feline kidney are significantly higher than that of other assessed tissues, and viral infection resulted in a significant upregulation of ACE2 RNA by 4 dpi and then a subsequent reduction by 8 dpi. Hypertension and activation of the renin-angiotensin system may have driven this rise in ACE2 in order to counterbalance system effects on infection, and future studies should include blood pressure evaluation in conjunction with other clinical parameters such as oxygen saturation, chemistry panels, and imaging. Further studies are needed to fully understand the role of ACE2 in SARS-CoV-2 viral replication kinetics and disease.

Limitations to this study include sample sizes as well as sampling time points. Further studies are needed to try and better identify peak viral loads in various tissues as well as ACE2 expression in order to investigate when viral clearance occurs and when this feline model moves from early exudative disease to more organized, fibrotic disease. A better delineation of these events would add value to the model and its potential for use at other stages of disease. In addition, transmission and viral shedding after intra-tracheal inoculation of SARS-CoV-2 should be evaluated and compared with that of intra-nasal inoculation and spread. Expansion of in depth diagnostics was limited due to animal biosafety level requirements and availability of resources, but future studies will seek to evaluate other clinical parameters, such as oxygen saturation, thoracic imaging and complete blood counts, chemistry panels and urine analysis to better assess damage to other organ systems and further compare with human disease.

This feline model of SARS-CoV-2 infection offers an animal model that closely mirrors both clinical disease and pathology identified in hospitalized patients with severe COVID-19, making the model a potential option for future studies addressing novel therapeutics for COVID-19. Therapeutic measures can be thoroughly assessed for improvement in pathology and mitigation of clinical disease in cats before being validated in human trials, and more thorough evaluation of the feline immune response to infection may elucidate other options for COVID-19 treatments that could mitigate disease and improve clinical outcomes. The continued emergence of novel variants, circulating globally, ceaselessly contributes to the complexity and duration of this pandemic. This animal model offers an ease of use, which can positively impact further vaccination and control strategies necessary to achieve an end to the rapid spread of COVID-19. This model also offers utility in a One Health approach to the role of companion animals in disease transmission, antigenic drift, and more thorough evaluation of the potential for feline contributions to the spread of SARS-CoV-2.

## MATERIALS AND METHODS

### Ethics Statement

This study was approved by the Oklahoma State University Institutional Animal Care and Use Committee; IACUC-20-48, Validation of a naturally occurring animal model for SARS-CoV-2 infection. Oklahoma State University’s animal care and use program is licensed by the United States Department of Agriculture (USDA), and accredited by the Association for Assessment and Accreditation of Laboratory Animal Care (AAALAC). In accordance with the approved IACUC protocol, animals were monitored at least once daily for evidence of morbidity and discomfort by trained animal care staff. Prior to experimental procedures, all study animals were anesthetized to minimize animal suffering and distress. Human euthanasia procedures were conducted by phenobarbital overdose in accordance with IACUC protocols and American Veterinary Medical Association (AVMA) Guidelines for the Euthanasia of Animals. Prior to euthanasia, all study animals were anesthetized by intramuscular injection of ketamine (4 mg/kg), dexmedetomidine (0.02 mg/kg), and butorphanol (0.4 mg/kg). No animals died without euthanasia during this study.

### Virus

SARS-CoV-2 virus isolate USA-WA1/2020 was obtained from BEI Resources, passaged up to 6 times in Vero E6 cells in Vero E6 cell growth medium. Virus stock was titrated and quantified on Vero E6 cells using a standard MERS-CoV quantification assay (45). TCID_50_ was calculated using the Reed and Muench method.

### Animals

Eighteen adult (9 males, 9 females, all 9 months-old) specific pathogen free (SPF) cats were obtained from Marshall BioResources (North Rose, NY). Animals intended for SARS-CoV-2 inoculation were housed within Biosafety Level 3 (BSL-3) barrier animal rooms at Oklahoma State University, individually housed and fed dry/wet food with access to water *ad libitum*. Animals intended for sham-inoculation were group-housed within the AAALAC International Accredited animal facility at Oklahoma State University. Animals were allowed 30 days for acclimation prior to initiation of the study. Temperature-sensing microchips (Bio Medic Data Systems, Seaford, DE) were implanted subcutaneously in the dorsum after 30 days.

Baseline weights, body temperatures, clinical evaluation, and nasal swab sampling were obtained prior to inoculation. All animals were in apparent good health at the onset of the study.

### Virus Challenge

Cats were lightly anesthetized with ketamine (4 mg/kg), dexmedetomidine (20 µg/kg), and butorphanol (0.4 mg/kg) intramuscularly. Cats were then positioned in ventral recumbency and intubated so that the end of an endotracheal tube is positioned within the distal trachea as described (46). In twelve cats, a 3-cc syringe was used to inoculate 1 mL 9⨯10^5^ PFU (1.26⨯10^6^ TCID_50_) per mL SARS-CoV-2, isolate USA-WA1/2020 in Dulbecco’s Modified Eagle Medium (DMEM), followed by 2 mL of air from an empty syringe. The remaining six cats were sham-inoculated using sterile PBS via the same method. Viral inoculum dosage was confirmed through virus back-titration on E6 cells immediately following inoculation.

### Sampling

Blood and nasal swab samples were collected under sedation (described above) from all study animals (n=18) at day 0 to serve as baseline for ddPCR analysis. Blood samples (6 mL) were obtained from all cats via cephalic or medial saphenous venipuncture and processed immediately for viral quantification. Nasal swab samples obtained from the nares of all cats using ultrafine flocked swabs (Puritan) were placed in 2 mL microcentrifuge tubes containing RNAlater solution (Ambio, Austin, TX) and stored at −80°C until processed. At day 4 and day 8 post-inoculation, a subset of SARS-CoV-2 infected cats (n=6 per time point) and sham-inoculated control cats (n=3 per time point) were anesthetized for blood and nasal swab collection then humanely euthanized (pentobarbital >80mg/kg) and necropsied to collect tissue samples. Necropsied tissues were processed for histologic examination, immunohistochemistry (IHC), and RNA analysis as described below.

### Clinical Observations and Scoring

Animals were monitored at least once daily for evidence of morbidity and discomfort by a licensed veterinary practitioner. Body weights and temperatures (thermal microchips) were documented daily every morning for the duration of the study. Full clinical scoring included evaluation of body weight, body temperature, activity levels, behavior, respiratory effort, evidence of ocular/nasal discharge, and recognition of coughing or wheezing. Each factor was assigned a score of 0 (normal), 1 (mild-moderate), or 2 (severe) as described in Table 1. Each clinical factor parameter was added to assign an individual animal a summed clinical score every 24 hours for the duration of the study. Cats were observed at rest for respiration rates, activity levels and other notable clinical signs before stimulation.

### Histopathology

Necropsy was performed on six (n=6) SARS-CoV-2-infected cats at 4 dpi and the remaining six (n=6) SARS-CoV-2-infected cats at 8 dpi. Three (n=3) sham-inoculated cats were necropsied at each time point (4 dpi and 8 dpi) to provide control samples. Tissue collection included: lung, tracheobronchial lymph nodes, nasal turbinates, distal trachea, and kidney. Necropsy tissues were halved and then placed into either 1 mL tubes and frozen at − 80°C, or into standard tissue cassettes that were then fixed in 10% neutral-buffered formaldehyde for 96 hours prior transferring to 70% ethanol for 72 hours. Tissues were then trimmed and processed for histology. Five µm paraffin sections were collected onto charged slides, and one slide of each tissue was stained with hematoxylin and eosin (H & E) for microscopic evaluation. Necropsy tissues were evaluated for evidence of inflammation and/or aberrations in lymphoid populations as reported in human COVID-19 patients (34-37). Lung tissues were specifically evaluated for the following pathology: alveolar damage (pneumocyte necrosis, hyaline membrane formation,) alveolar fibrin deposition (± organization), serous exudate/edema, perivascular infiltrates, alveolar histiocytes, type II pneumocyte hyperplasia, syncytia, thrombosis, and fibrinoid vasculitis. All tissues were assigned a quantitative histologic score based on previously documented criteria (47, 48): 0 = no apparent pathology/change; 1 = minimal change (minimally increased numbers of inflammatory cells); 2 = mild change (mild inflammatory infiltrates, alveolar damage/necrosis, fibrin deposition and/or exudation); 3 = moderate change (as previously described, but more moderately extensive); 4 = marked changes (as previously described, but with severe inflammation, alveolar damage, hyaline membrane formation, necrosis, exudation, vasculitis and/or thrombosis). All tissues were evaluated and scored by a board-certified veterinary pathologist blinded to study groups to ensure scientific rigor and reproducibility.

### Viral RNA Analysis

Viral RNA analysis was performed on samples from nasal swabs, collected plasma, and tissues. Nasal swabs were immediately broken off into 1.5 mL microcentrifuge tubes containing 200 µL RNAlater Solution (Ambion, Austin, TX) and stored at −20°C. The nasal swabs were vortexed for 15 seconds, then inverted and centrifuged at 1500 rpm for 10 minutes. RNA was extracted from frozen necropsy tissues using a QIAamp Viral RNA Mini Kit (Qiagen, Germantown, MD) and tissue homogenizer. SARS-CoV-2 viral RNA was quantified by droplet digital PCR (ddPCR) as previously described (49). Briefly, ddPCR was performed according to manufacturer’s instructions for the 2019-nCoV CDC ddPCR Triplex Probe Assay (Bio-Rad, Hercules, California, USA). PCR reaction mixtures were as follows: 5.5 μl One-Step RT-ddPCR Advanced Kit for Probes Supermix (no dUTP’s) (Bio-Rad), 2.2 μl reverse transcriptase, 1.1 μl 300 mM Dithiothreitol (DTT), 1.1 μl triplex probe assay (for N1, N2, RPP30 detection), 2.2 μl RNase free water, and 9.9 μl RNA template in a final volume of 22 μl per sample. Duplicate 20 μl samples were partitioned using a QX200 droplet generator (Bio-Rad, Hercules, California, USA) and then transferred to a 96-well plate and sealed. Samples were processed in a C1000 touch Thermal Cycler (Bio-Rad) under the following cycling protocol: 50 °C for 60 min for reverse transcription, 95 °C for 10 min for enzyme activation, 94°C for 30 s for denaturation and 55 °C for 60 s for annealing/extension for 45 cycles, 98 °C 10 min for enzyme deactivation, 4 °C for 30 min for droplet stabilization followed by infinite 4 °C hold. The amplified samples were read in the FAM and HEX channels using the QX200 reader (Bio-Rad). Each experiment was performed with a negative control (no template control, NTC) and a positive control (RNA extracted from SARS-CoV-2 viral stock and diluted 1:12,000). Data were analyzed using QuantaSoft™ v1 AnalysisPro Software (Bio-Rad) and expressed as Log_10_ (copies/mL).

### Feline ACE2 Analysis

Feline angiotensin converting enzyme 2 (fACE2) RNA was quantified by ddPCR using methods similar to the above assay for CoV. RNA was extracted from frozen necropsy tissues as outlined above. cDNA was synthesized as previously published (48). Design of primers and probe targeting fACE2 was performed according to manufacturer’s recommendation, namely keeping GC content between 50–60 % for primers and 30–80 % for probes, and melting temperatures between 50–65 °C for primers and 3–10 °C higher for probes. Oligoes were synthesized by Integrated DNA Technologies (IDT, Coralville, Iowa, USA). The sequences are as follows: Forward: 5′-ACGGAGGCGTAAGGATTT-3′, Reverse: 5′ - GTGTGGTAGTGGTTGGTATTG-3′, probe: 5′ - CGGGATCAGAAATCGAAGGAAGAA - 3′. BLAST analysis (50) of the primer and probe sequences against the domestic cat (*Felis catus*) genome was performed to ensure no similar sequences could be amplified. ddPCR reactions were prepared by adding 11 μl Supermix for Probes (no dUTP) (Bio-Rad), 1.1 μl of primer/probe mix (final concentration is 500nM for primers and 250 nM for probe) and 8.8 μl of cDNA template containing 110 ng RNA equivalent. Droplets were partitioned and PCR executed as above using the following cycling conditions: 95 °C for 10 min, 95 °C 30 s for denaturation and 58.8 °C for 60 s for annealing/extension for 45 cycles, 98 °C 10 min for enzyme deactivation. Droplets were read and analyzed as described above.

### Immunohistochemistry

5µm sections of formalin-fixed, paraffin-embedded lung were mounted on charged glass slides, baked for one hour at 60°C, and passed through Xylene, graded ethanol, and double distilled water to remove paraffin and rehydrate tissue sections. A microwave was used for heat induced epitope retrieval. Slides were heated in a high pH solution (Vector Labs H-3301), rinsed in hot water and transferred to a heated low pH solution (Vector Labs H-3300) where they were allowed to cool to room temperature. Sections were washed in a solution of phosphate-buffered saline and fish gelatin (PBS-FSG) and transferred to a humidified chamber, for staining at room temperature. Tissues were blocked with 10% normal goat serum (NGS) for 40 minutes, followed by a 60-minute incubation with a guinea pig anti-SARS antibody (BEI NR-10361) diluted 1:1000 in NGS. Slides were washed and transferred to the humidified chamber for a 40-minute incubation with a goat anti-guinea pig secondary antibody (Invitrogen A11073) tagged with Alexa Fluor 488 and diluted 1:1000 in NGS. Following washes, DAPI (4’,6-diamidino-2-phenylindole) was used to label the nuclei of each section. Slides were mounted using a homemade anti-quenching mounting media containing Mowiol (Calbiochem#475904) and DABCO (Sigma #D2522) and imaged at 20X with a Zeiss Axio Slide Scanner.

### Statistical Analyses

When applicable, data were expressed as mean ± SEM and statistically analyzed using GraphPad Prism 9.0 software (La Jolla, CA). Kruskal–Wallis test, Pearson correlations, and ANOVA were used to compare differences in clinical score, histopathology, SARS-CoV-2 viral load, and ACE2 RNA among uninfected and SARS-CoV-2-infected individuals, between sample type, for each tissue individually, and between tissues. For all significant results, pair-wise comparisons were made by post-hoc analysis. P-values < 0.05 were considered significant.

## Acknowledgments

The authors would like to thank Girish Patil, Akhilesh Ramachandran, and Sai Narayanan with the Oklahoma Animal Disease Diagnostic Lab for their assistance in viral RNA extraction and technical advisement. We would also like to acknowledge Curtis Andrew for assistance with slide preparation and coordination of immunohistochemistry from the Immunopathology Core Laboratory at Oklahoma State College of Veterinary Medicine.

## Author Contributions

Conceptualization: CAM, JMR, JWR

Methodology: CAM, JMR, JWR, SC, CCM

Investigation: CAM, JMR, MTS, SC, JWR, CCM

Visualization: CAM, JMR, MTS

Funding Acquisition: CAM

Writing – original draft: JMR, CAM

Writing – review and editing: JWR, CCM, SC, MTS

## Data and materials availability

All relevant data associated with this study has been deposited in a public repository: 10.6084/m9.figshare.14449773

## References

1. WHO. World Health Organization Coronavirus (COVID-19) Dashboard: World Health Organization; 2021 [cited 2021. Available from: https://covid19.who.int.

2. Wu C, Chen X, Cai Y, Zhou X, Xu S, Huang H, et al. Risk factors associated with acute respiratory distress syndrome and death in patients with coronavirus disease 2019 pneumonia in Wuhan, China. JAMA internal medicine. 2020.

3. Chen G, Wu D, Guo W, Cao Y, Huang D, Wang H, et al. Clinical and immunologic features in severe and moderate Coronavirus Disease 2019. The Journal of Clinical Investigation. 2020.

4. Pedersen SF, Ho Y-C. SARS-CoV-2: a storm is raging. The Journal of Clinical Investigation. 2020.

5. Kumar S, Yadav PK, Srinivasan R, Perumal N. Selection of animal models for COVID-19 research. Virusdisease. 2020;31(4):1–6.

6. Gretebeck LM, Subbarao K. Animal models for SARS and MERS coronaviruses. Curr Opin Virol. 2015;13:123–9.

7. Hikmet F, Méar L, Edvinsson Å, Micke P, Uhlén M, Lindskog C. The protein expression profile of ACE2 in human tissues. Mol Syst Biol. 2020;16(7):e9610.

8. Kiros M, Andualem H, Kiros T, Hailemichael W, Getu S, Geteneh A, et al. COVID-19 pandemic: current knowledge about the role of pets and other animals in disease transmission. Virol J. 2020;17(1):143.

9. Oreshkova N, Molenaar RJ, Vreman S, Harders F, Oude Munnink BB, Hakze-van der Honing RW, et al. SARS-CoV-2 infection in farmed minks, the Netherlands, April and May 2020. Euro Surveill. 2020;25(23).

10. Newman A, Smith D, Ghai RR, Wallace RM, Torchetti MK, Loiacono C, et al. First Reported Cases of SARS-CoV-2 Infection in Companion Animals - New York, March-April 2020. MMWR Morb Mortal Wkly Rep. 2020;69(23):710–3.

11. Molenaar RJ, Vreman S, Hakze-van der Honing RW, Zwart R, de Rond J, Weesendorp E, et al. Clinical and Pathological Findings in SARS-CoV-2 Disease Outbreaks in Farmed Mink (Neovison vison). Vet Pathol. 2020;57(5):653–7.

12. Luan J, Lu Y, Jin X, Zhang L. Spike protein recognition of mammalian ACE2 predicts the host range and an optimized ACE2 for SARS-CoV-2 infection. Biochem Biophys Res Commun. 2020;526(1):165–9.

13. Liu Y, Hu G, Wang Y, Ren W, Zhao X, Ji F, et al. Functional and genetic analysis of viral receptor ACE2 orthologs reveals a broad potential host range of SARS-CoV-2. Proc Natl Acad Sci U S A. 2021;118(12).

14. Hayashi T, Abiko K, Mandai M, Yaegashi N, Konishi I. Highly conserved binding region of ACE2 as a receptor for SARS-CoV-2 between humans and mammals. Vet Q. 2020;40(1):243–9.

15. McAloose D, Laverack M, Wang L, Killian ML, Caserta LC, Yuan F, et al. From People to Panthera: Natural SARS-CoV-2 Infection in Tigers and Lions at the Bronx Zoo. mBio. 2020;11(5).

16. Shi J, Wen Z, Zhong G, Yang H, Wang C, Huang B, et al. Susceptibility of ferrets, cats, dogs, and other domesticated animals to SARS-coronavirus 2. Science. 2020.

17. Zhang Q, Zhang H, Gao J, Huang K, Yang Y, Hui X, et al. A serological survey of SARS-CoV-2 in cat in Wuhan. Emerg Microbes Infect. 2020;9(1):2013–9.

18. Lakdawala SS, Menachery VD. The search for a COVID-19 animal model. Science. 2020;368(6494):942–3.

19. Hernández M, Abad D, Eiros JM, Rodríguez-Lázaro D. Are Animals a Neglected Transmission Route of SARS-CoV-2? Pathogens. 2020;9(6).

20. Van de Velde H, Janssens GP, de Rooster H, Polis I, Peters I, Ducatelle R, et al. The cat as a model for human obesity: insights into depot-specific inflammation associated with feline obesity. Br J Nutr. 2013;110(7):1326–35.

21. Nelson RW, Reusch CE. Animal models of disease: classification and etiology of diabetes in dogs and cats. J Endocrinol. 2014;222(3):T1–9.

22. Samaha G, Beatty J, Wade CM, Haase B. The Burmese cat as a genetic model of type 2 diabetes in humans. Anim Genet. 2019;50(4):319–25.

23. Hoenig M. The cat as a model for human obesity and diabetes. J Diabetes Sci Technol. 2012;6(3):525–33.

24. Wallner M, Eaton DM, Berretta RM, Borghetti G, Wu J, Baker ST, et al. A Feline HFpEF Model with Pulmonary Hypertension and Compromised Pulmonary Function. Sci Rep. 2017;7(1):16587.

25. Prat V, Rozec B, Gauthier C, Lauzier B. Human heart failure with preserved ejection versus feline cardiomyopathy: what can we learn from both veterinary and human medicine? Heart Fail Rev. 2017;22(6):783–94.

26. Gaudreault NN, Trujillo JD, Carossino M, Meekins DA, Morozov I, Madden DW, et al. SARS-CoV-2 infection, disease and transmission in domestic cats. Emerg Microbes Infect. 2020;9(1):2322–32.

27. Bosco-Lauth AM, Hartwig AE, Porter SM, Gordy PW, Nehring M, Byas AD, et al. Experimental infection of domestic dogs and cats with SARS-CoV-2: Pathogenesis, transmission, and response to reexposure in cats. Proc Natl Acad Sci U S A. 2020;117(42):26382–8.

28. Gaudreault NN, Carossino M, Morozov I, Trujillo JD, Meekins DA, Madden DW, et al. Experimental re-infected cats do not transmit SARS-CoV-2. Emerg Microbes Infect. 2021;10(1):638–50.

29. van den Brand JM, Haagmans BL, Leijten L, van Riel D, Martina BE, Osterhaus AD, et al. Pathology of experimental SARS coronavirus infection in cats and ferrets. Vet Pathol. 2008;45(4):551–62.

30. van den Brand JM, Haagmans BL, van Riel D, Osterhaus AD, Kuiken T. The pathology and pathogenesis of experimental severe acute respiratory syndrome and influenza in animal models. J Comp Pathol. 2014;151(1):83–112.

31. Huet O, Ramsey D, Miljavec S, Jenney A, Aubron C, Aprico A, et al. Ensuring animal welfare while meeting scientific aims using a murine pneumonia model of septic shock. Shock. 2013;39(6):488–94.

32. Hartmann AD, Helps CR, Lappin MR, Werckenthin C, Hartmann K. Efficacy of pradofloxacin in cats with feline upper respiratory tract disease due to Chlamydophila felis or Mycoplasma infections. J Vet Intern Med. 2008;22(1):44–52.

33. Steagall PV, Monteiro BP. Acute pain in cats: Recent advances in clinical assessment. J Feline Med Surg. 2019;21(1):25–34.

34. Polak SB, Van Gool IC, Cohen D, Jan H, van Paassen J. A systematic review of pathological findings in COVID-19: a pathophysiological timeline and possible mechanisms of disease progression. Modern Pathology. 2020;33(11):2128–38.

35. Tian S, Xiong Y, Liu H, Niu L, Guo J, Liao M, et al. Pathological study of the 2019 novel coronavirus disease (COVID-19) through postmortem core biopsies. Modern Pathology. 2020;33(6):1007–14.

36. von der Thüsen J, van der Eerden M. Histopathology and genetic susceptibility in COVID-19 pneumonia. European journal of clinical investigation. 2020;50(7):e13259.

37. Xu Z, Shi L, Wang Y, Zhang J, Huang L, Zhang C, et al. Pathological findings of COVID-19 associated with acute respiratory distress syndrome. The Lancet respiratory medicine. 2020;8(4):420–2.

38. Bodewes R, Kreijtz JH, van Amerongen G, Fouchier RA, Osterhaus AD, Rimmelzwaan GF, et al. Pathogenesis of Influenza A/H5N1 virus infection in ferrets differs between intranasal and intratracheal routes of inoculation. Am J Pathol. 2011;179(1):30–6.

39. Rimmelzwaan GF, van Riel D, Baars M, Bestebroer TM, van Amerongen G, Fouchier RA, et al. Influenza A virus (H5N1) infection in cats causes systemic disease with potential novel routes of virus spread within and between hosts. Am J Pathol. 2006;168(1):176-83; quiz 364.

40. Macdonald ES, Norris CR, Berghaus RB, Griffey SM. Clinicopathologic and radiographic features and etiologic agents in cats with histologically confirmed infectious pneumonia: 39 cases (1991-2000). J Am Vet Med Assoc. 2003;223(8):1142–50.

41. Wiersinga WJ, Rhodes A, Cheng AC, Peacock SJ, Prescott HC. Pathophysiology, Transmission, Diagnosis, and Treatment of Coronavirus Disease 2019 (COVID-19): A Review. Jama. 2020;324(8):782–93.

42. Xu Z, Shi L, Wang Y, Zhang J, Huang L, Zhang C, et al. Pathological findings of COVID-19 associated with acute respiratory distress syndrome. Lancet Respir Med. 2020;8(4):420–2.

43. Fox SE, Akmatbekov A, Harbert JL, Li G, Quincy Brown J, Vander Heide RS. Pulmonary and cardiac pathology in African American patients with COVID-19: an autopsy series from New Orleans. Lancet Respir Med. 2020;8(7):681–6.

44. Borczuk AC, Salvatore SP, Seshan SV, Patel SS, Bussel JB, Mostyka M, et al. COVID-19 pulmonary pathology: a multi-institutional autopsy cohort from Italy and New York City. Mod Pathol. 2020;33(11):2156–68.

45. Coleman CM, Frieman MB. Growth and quantification of MERS-CoV infection. Current protocols in microbiology. 2015;37(1):15E. 2.1-E. 2.9.

46. Ritchey JW, Levy JK, Bliss SK, Tompkins WA, Tompkins MB. Constitutive expression of types 1 and 2 cytokines by alveolar macrophages from feline immunodeficiency virus-infected cats. Vet Immunol Immunopathol. 2001;79(1-2):83–100.

47. Miller C, Bielefeldt-Ohmann H, MacMillan M, Huitron-Resendiz S, Henriksen S, Elder J, et al. Strain-specific viral distribution and neuropathology of feline immunodeficiency virus. Veterinary immunology and immunopathology. 2011;143(3):282–91.

48. Miller C, Boegler K, Carver S, MacMillan M, Bielefeldt-Ohmann H, VandeWoude S. Pathogenesis of oral FIV infection. PloS one. 2017;12(9):e0185138.

49. Deiana M, Mori A, Piubelli C, Scarso S, Favarato M, Pomari E. Assessment of the direct quantitation of SARS-CoV-2 by droplet digital PCR. Scientific reports. 2020;10(1):1–7.

50. O’Driscoll A, Belogrudov V, Carroll J, Kropp K, Walsh P, Ghazal P, et al. HBLAST: Parallelised sequence similarity--A Hadoop MapReducable basic local alignment search tool. J Biomed Inform. 2015;54:58–64.

51. Blair RV, Vaccari M, Doyle-Meyers LA, Roy CJ, Russell-Lodrigue K, Fahlberg M, et al. Acute Respiratory Distress in Aged, SARS-CoV-2–Infected African Green Monkeys but Not Rhesus Macaques. The American journal of pathology. 2021;191(2):274–82.

