## Supplementary material for "Clinicopathologic features of a feline SARS-CoV-2 infection model parallel acute COVID-19 in humans": S1 Table, S2 Table, S3 Table, S4 Table

**S1 Table. Clinical Scoring Data Summary.** (A) Total number of cats from each group exhibiting clinical signs during the 8-day study. (B) Total clinical scores recorded each day per individual cat. Statistical analysis and raw scoring data for each day is included in Supporting Information.

**A.**

**Total # Cats With Specific Clinical Sign During Study**

| **Clinical Sign** | **PBS (n=6)** | **SARS-CoV-2 (n=12)** |
|---|---|---|
| Weight loss | 1 | 5 |
| Reduced activity | 0 | 12 |
| Altered behavior | 0 | 7 |
| Hyperthermia | 0 | 8 |
| Tachypnea or Dyspnea | 0 | 12 |
| Ocular or Nasal Discharge | 0 | 0 |
| Coughing/Wheezing | 0 | 4 |

**B.**

**Total Clinical Score (Sum of parameter scores)**

| | Day 0 | Day 1 | Day 2 | Day 3 | Day 4 | Day 5 | Day 6 | Day 7 | Day 8 | **Peak Clinical Score** |
|---|---|---|---|---|---|---|---|---|---|---|
| Control 1 (0109) | 0 | 0 | 0 | 0 | 0 | | | | | **0** |
| Control 2 (0575) | 0 | 0 | 0 | 0 | 0 | | | | | **0** |
| Control 3 (0664) | 0 | 0 | 0 | 0 | 0 | | | | | **0** |
| Control 4 (1024) | 0 | 0 | 0 | 0 | 0 | 0 | 0 | 0 | 0 | **0** |
| Control 5 (0923) | 0 | 0 | 0 | 0 | 0 | 0 | 0 | 0 | 0 | **0** |
| Control 6 (0907) | 0 | 0 | 0 | 0 | 0 | 1 | 0 | 0 | 0 | **0** |
| Infected 1 (9491) | 0 | 0 | 0 | 0 | 2 | | | | | **2** |
| Infected 2 (0613) | 0 | 3 | 3 | 2 | 2 | | | | | **3** |
| Infected 3 (9687) | 0 | 0 | 0 | 0 | 2 | | | | | **2** |
| Infected 4 (9857) | 0 | 0 | 0 | 0 | 2 | | | | | **2** |
| Infected 5 (0452) | 0 | 0 | 2 | 7 | 7 | | | | | **7** |
| Infected 6 (1474) | 0 | 1 | 5 | 3 | 6 | | | | | **6** |
| Infected 7 (1440) | 0 | 1 | 0 | 0 | 6 | 4 | 2 | 2 | 3 | **6** |
| Infected 8 (0800) | 0 | 2 | 0 | 0 | 2 | 4 | 2 | 3 | 2 | **4** |
| Infected 9 (0559) | 0 | 2 | 2 | 1 | 9 | 4 | 3 | 6 | 6 | **9** |
| Infected 10 (1130) | 0 | 0 | 0 | 3 | 2 | 5 | 2 | 2 | 2 | **5** |
| Infected 11 (0095) | 0 | 0 | 1 | 0 | 2 | 1 | 2 | 2 | 3 | **3** |
| Infected 12 (0842) | 0 | 1 | 1 | 0 | 2 | 2 | 2 | 2 | 2 | **2** |

**S2 Table.** Lung tissue histopathology scores based on pathologic features reported in human COVID-19 patients [33-36]. All tissues were assigned a quantitative histologic score of 0-4 based on previously documented criteria [37,38]

| | **Cat ID** | Alveolar Damage | Alveolar fibrin | Serous exudate/ pulmonary edema | Perivascular Infiltrates | Alveolar Histiocytosis | Type II Pneumocyte Hyperplasia | Syncytia | Thrombosis | Vasculitis | **Total Histopathology Score** |
|---|---|---|---|---|---|---|---|---|---|---|---|
| Uninfected | 0109 | 0 | 0 | 1 | 1 | 1 | 0 | 0 | 0 | 0 | **3** |
| | 0575 | 0 | 0 | 0 | 0 | 1 | 0 | 0 | 0 | 0 | **1** |
| | 0664 | 0 | 0 | 1 | 1 | 0 | 0 | 0 | 0 | 0 | **2** |
| | 0907 | 0 | 0 | 1 | 0 | 1 | 0 | 0 | 0 | 0 | **2** |
| | 0923 | 0 | 0 | 1 | 2 | 2 | 1 | 0 | 0 | 0 | **6** |
| | 1024 | 0 | 0 | 0 | 1 | 1 | 0 | 0 | 0 | 0 | **2** |
| Day 4 p.i. | 0452 | 3 | 3 | 4 | 4 | 4 | 3 | 2 | 0 | 0 | **23** |
| | 0613 | 0 | 0 | 2 | 2 | 0 | 0 | 0 | 0 | 0 | **4** |
| | 1474 | 2 | 0 | 0 | 3 | 4 | 2 | 2 | 0 | 0 | **13** |
| | 9491 | 2 | 0 | 4 | 3 | 3 | 0 | 0 | 0 | 0 | **12** |
| | 9687 | 2 | 0 | 1 | 3 | 3 | 0 | 0 | 0 | 0 | **9** |
| | 9857 | 3 | 2 | 3 | 3 | 4 | 0 | 0 | 0 | 0 | **15** |
| Day 8 p.i. | 0095 | 1 | 0 | 2 | 3 | 3 | 0 | 0 | 0 | 0 | **9** |
| | 0559 | 2 | 2 | 4 | 3 | 4 | 0 | 0 | 2 | 1 | **17** |
| | 0800 | 0 | 0 | 1 | 1 | 1 | 0 | 0 | 2 | 0 | **5** |
| | 0842 | 0 | 0 | 2 | 1 | 2 | 0 | 0 | 0 | 0 | **5** |
| | 1130 | 3 | 2 | 4 | 4 | 4 | 3 | 3 | 0 | 2 | **23** |
| | 1440 | 1 | 1 | 3 | 3 | 3 | 0 | 0 | 0 | 0 | **11** |

**S3 Table.** Total SARS-CoV-2 viral RNA and fACE2 RNA (copy number/ml) detected in each tissue type.

| | | CoV-2 Viral RNA Copies/ml | | | | | ACE2 RNA Copies/ml | | | | |
|---|---|---|---|---|---|---|---|---|---|---|---|
| | **Cat ID** | Nasal Turbinates | TB LN | Distal Trachea | Kidney | Lung | Nasal Turbinates | TB LN | Distal Trachea | Kidney | Lung |
| Uninfected | 0109 | 0 | 0 | 0 | 0 | 0 | 4178 | 0 | 1511 | 42667 | 133 |
| | 0575 | 0 | 0 | 0 | 0 | 0 | 138889 | 0 | 133 | 32667 | 778 |
| | 0664 | 0 | 0 | 0 | 0 | 0 | 3956 | 0 | 1711 | 189333 | 0 |
| | 0907 | 0 | 0 | 0 | 0 | 0 | 3200 | 311 | 0 | 192000 | 578 |
| | 0923 | 0 | 0 | 0 | 0 | 0 | 1578 | 1111 | 533 | 147778 | 0 |
| | 1024 | 0 | 0 | 0 | 0 | 0 | 4489 | 0 | 133 | 51778 | 0 |
| Day 4 p.i. | 0452 | 2755 | 214666 | 7333 | 267 | 488667 | 778 | 178 | 0 | 242444 | 0 |
| | 0613 | 378 | 244 | 14444 | 3178 | 3778 | 667 | 0 | 133 | 263111 | 133 |
| | 1474 | 1622 | 524444 | 178 | 1400 | 594889 | 1289 | 0 | 467 | 189556 | 533 |
| | 9491 | 104667 | 1889 | 0 | 622 | 1489 | 733 | 0 | 133 | 282000 | 133 |
| | 9687 | 16000 | 253555 | 2445 | 1644 | 866 | 64889 | 0 | 400 | 833333 | 0 |
| | 9857 | 47111 | 8889 | 4334 | 489 | 503778 | 25333 | 267 | 0 | 206667 | 0 |
| Day 8 p.i. | 0095 | 0 | 226666 | 32000 | 1755 | 1133 | 2644 | 422 | 133 | 148222 | 0 |
| | 0559 | 178 | 384445 | 200 | 222 | 28666 | 2356 | 0 | 0 | 150667 | 0 |
| | 0800 | 712 | 22667 | 63778 | 1711 | 8000 | 8222 | 10444 | 267 | 129333 | 667 |
| | 0842 | 956 | 189556 | 0 | 2889 | 778 | 156 | 2600 | 133 | 65556 | 0 |
| | 1130 | 11111 | 334000 | 600 | 1889 | 9267 | 1000 | 0 | 267 | 76222 | 133 |
| | 1440 | 3755 | 768889 | 24000 | 644 | 6444 | 1822 | 133 | 0 | 58444 | 0 |

* TB LN = tracheobronchial lymph node.

**S4 Table.** Relationship between feline ACE2 expression and tissue type (Kruskal-Wallis test). (A) Sham-inoculated control animals; (B) SARS-CoV-2-infected animals, 4 dpi; (C) SARS-CoV-2-infected animals, 8 dpi.

**A. Uninfected control animals (n=6): ACE2 RNA expression by tissue type**

| **Kruskal-Wallis Test** | Mean rank diff. | Significant? | Summary | P value | |
|---|---|---|---|---|---|
| Nasal Turbinates vs. TbLn | 14.17 | Yes | ** | 0.0049 | A-B |
| Nasal Turbinates vs. Distal Trachea | 9.583 | No | ns | 0.0569 | A-C |
| Nasal Turbinates vs. Kidney | -5.167 | No | ns | 0.3046 | A-D |
| Nasal Turbinates vs. Lung | 13.08 | Yes | ** | 0.0093 | A-E |
| TbLn vs. Distal Trachea | -4.583 | No | ns | 0.3624 | B-C |
| TbLn vs. Kidney | -19.33 | Yes | *** | 0.0001 | B-D |
| TbLn vs. Lung | -1.083 | No | ns | 0.8296 | B-E |
| Distal Trachea vs. Kidney | -14.75 | Yes | ** | 0.0034 | C-D |
| Distal Trachea vs. Lung | 3.5 | No | ns | 0.4868 | C-E |
| Kidney vs. Lung | 18.25 | Yes | *** | 0.0003 | D-E |

| Test details | Mean rank 1 | Mean rank 2 | Mean rank diff. | n1 | n2 | Z |
|---|---|---|---|---|---|---|
| Nasal Turbinates vs. TbLn | 21.83 | 7.667 | 14.17 | 6 | 6 | 2.815 |
| Nasal Turbinates vs. Distal Trachea | 21.83 | 12.25 | 9.583 | 6 | 6 | 1.904 |
| Nasal Turbinates vs. Kidney | 21.83 | 27 | -5.167 | 6 | 6 | 1.027 |
| Nasal Turbinates vs. Lung | 21.83 | 8.75 | 13.08 | 6 | 6 | 2.6 |
| TbLn vs. Distal Trachea | 7.667 | 12.25 | -4.583 | 6 | 6 | 0.9107 |
| TbLn vs. Kidney | 7.667 | 27 | -19.33 | 6 | 6 | 3.842 |
| TbLn vs. Lung | 7.667 | 8.75 | -1.083 | 6 | 6 | 0.2153 |
| Distal Trachea vs. Kidney | 12.25 | 27 | -14.75 | 6 | 6 | 2.931 |
| Distal Trachea vs. Lung | 12.25 | 8.75 | 3.5 | 6 | 6 | 0.6955 |
| Kidney vs. Lung | 27 | 8.75 | 18.25 | 6 | 6 | 3.626 |

**B. SARS-CoV-2-infected animals, Day 4 p.i. (n=6): ACE2 RNA expression by tissue type**

| **Kruskal-Wallis Test** | Mean rank diff. | Significant? | Summary | P value | |
|---|---|---|---|---|---|
| Nasal Turbinates vs. TbLn | 13.33 | Yes | ** | 0.0078 | A-B |
| Nasal Turbinates vs. Distal Trachea | 10.5 | Yes | * | 0.036 | A-C |
| Nasal Turbinates vs. Kidney | -6 | No | ns | 0.2309 | A-D |
| Nasal Turbinates vs. Lung | 12.17 | Yes | * | 0.0151 | A-E |
| TbLn vs. Distal Trachea | -2.833 | No | ns | 0.5716 | B-C |
| TbLn vs. Kidney | -19.33 | Yes | *** | 0.0001 | B-D |
| TbLn vs. Lung | -1.167 | No | ns | 0.8158 | B-E |
| Distal Trachea vs. Kidney | -16.5 | Yes | *** | 0.001 | C-D |
| Distal Trachea vs. Lung | 1.667 | No | ns | 0.7393 | C-E |
| Kidney vs. Lung | 18.17 | Yes | *** | 0.0003 | D-E |

| Test details | Mean rank 1 | Mean rank 2 | Mean rank diff. | n1 | n2 | Z |
|---|---|---|---|---|---|---|
| Nasal Turbinates vs. TbLn | 21.5 | 8.167 | 13.33 | 6 | 6 | 2.662 |
| Nasal Turbinates vs. Distal Trachea | 21.5 | 11 | 10.5 | 6 | 6 | 2.096 |
| Nasal Turbinates vs. Kidney | 21.5 | 27.5 | -6 | 6 | 6 | 1.198 |
| Nasal Turbinates vs. Lung | 21.5 | 9.333 | 12.17 | 6 | 6 | 2.429 |
| TbLn vs. Distal Trachea | 8.167 | 11 | -2.833 | 6 | 6 | 0.5657 |
| TbLn vs. Kidney | 8.167 | 27.5 | -19.33 | 6 | 6 | 3.86 |
| TbLn vs. Lung | 8.167 | 9.333 | -1.167 | 6 | 6 | 0.2329 |
| Distal Trachea vs. Kidney | 11 | 27.5 | -16.5 | 6 | 6 | 3.294 |
| Distal Trachea vs. Lung | 11 | 9.333 | 1.667 | 6 | 6 | 0.3328 |
| Kidney vs. Lung | 27.5 | 9.333 | 18.17 | 6 | 6 | 3.627 |

**C. SARS-CoV-2-infected animals, Day 8 p.i. (n=6): ACE2 RNA expression by tissue type**

| **Kruskal-Wallis Test** | Mean rank diff. | Significant? | Summary | P value | |
|---|---|---|---|---|---|
| Nasal Turbinates vs. TbLn | 5.75 | No | ns | 0.2529 | A-B |
| Nasal Turbinates vs. Distal Trachea | 9.333 | No | ns | 0.0634 | A-C |
| Nasal Turbinates vs. Kidney | -8.333 | No | ns | 0.0975 | A-D |
| Nasal Turbinates vs. Lung | 11.58 | Yes | * | 0.0213 | A-E |
| TbLn vs. Distal Trachea | 3.583 | No | ns | 0.4761 | B-C |
| TbLn vs. Kidney | -14.08 | Yes | ** | 0.0051 | B-D |
| TbLn vs. Lung | 5.833 | No | ns | 0.246 | B-E |
| Distal Trachea vs. Kidney | -17.67 | Yes | *** | 0.0004 | C-D |
| Distal Trachea vs. Lung | 2.25 | No | ns | 0.6546 | C-E |
| Kidney vs. Lung | 19.92 | Yes | **** | <0.0001 | D-E |

| Test details | Mean rank 1 | Mean rank 2 | Mean rank diff. | n1 | n2 | Z |
|---|---|---|---|---|---|---|
| Nasal Turbinates vs. TbLn | 19.17 | 13.42 | 5.75 | 6 | 6 | 1.143 |
| Nasal Turbinates vs. Distal Trachea | 19.17 | 9.833 | 9.333 | 6 | 6 | 1.856 |
| Nasal Turbinates vs. Kidney | 19.17 | 27.5 | -8.333 | 6 | 6 | 1.657 |
| Nasal Turbinates vs. Lung | 19.17 | 7.583 | 11.58 | 6 | 6 | 2.303 |
| TbLn vs. Distal Trachea | 13.42 | 9.833 | 3.583 | 6 | 6 | 0.7126 |
| TbLn vs. Kidney | 13.42 | 27.5 | -14.08 | 6 | 6 | 2.801 |
| TbLn vs. Lung | 13.42 | 7.583 | 5.833 | 6 | 6 | 1.16 |
| Distal Trachea vs. Kidney | 9.833 | 27.5 | -17.67 | 6 | 6 | 3.513 |
| Distal Trachea vs. Lung | 9.833 | 7.583 | 2.25 | 6 | 6 | 0.4474 |
| Kidney vs. Lung | 27.5 | 7.583 | 19.92 | 6 | 6 | 3.961 |
