## Supplementary figures and images for "Clinicopathologic features of a feline SARS-CoV-2 infection model parallel acute COVID-19 in humans"

### S1 Fig

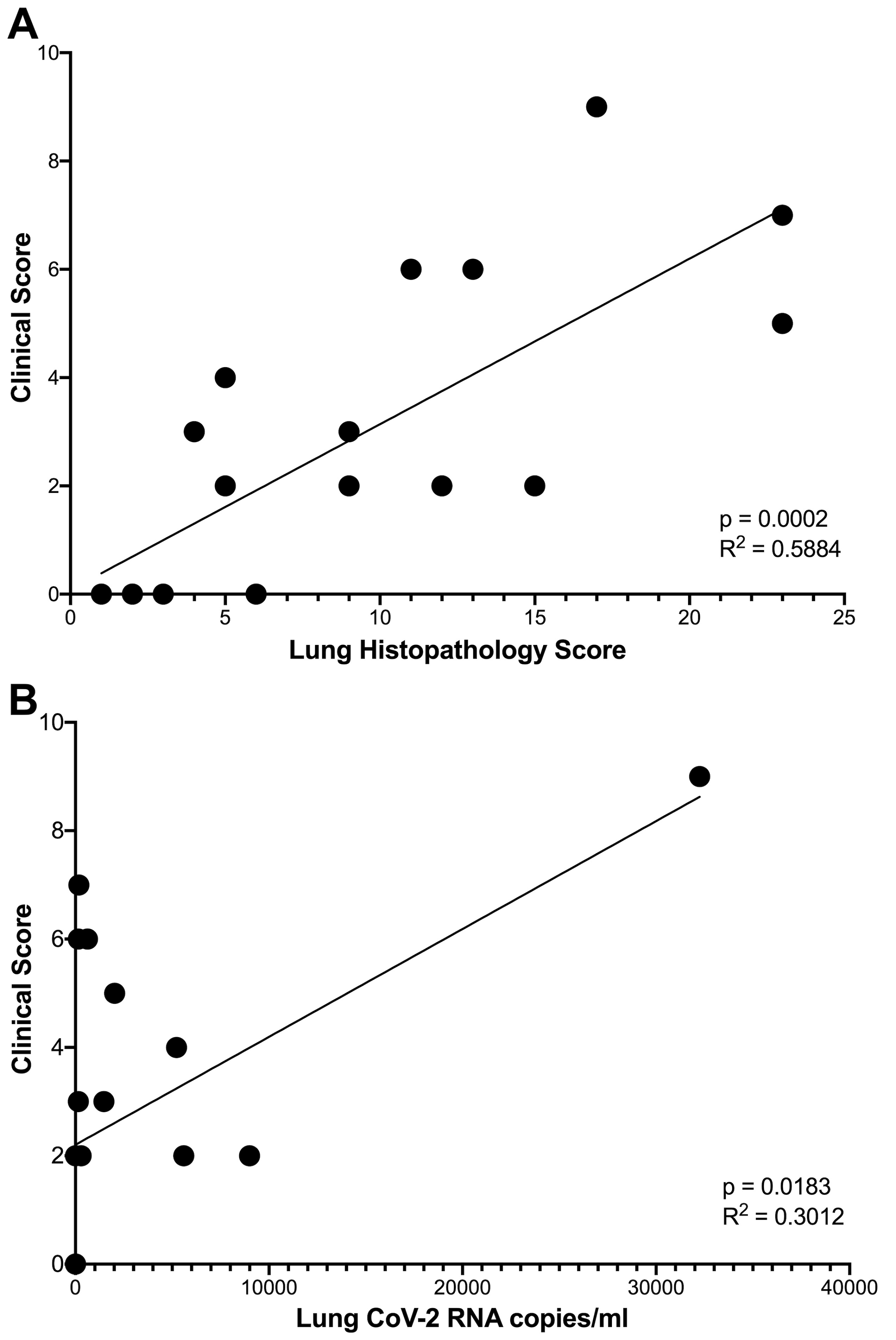
